# PrePCI: A structure- and chemical similarity-informed database of predicted protein compound interactions

**DOI:** 10.1101/2022.09.17.508184

**Authors:** Stephen J. Trudeau, Howook Hwang, Deepika Mathur, Kamrun Begum, Donald Petrey, Diana Murray, Barry Honig

## Abstract

We describe the Predicting Protein Compound Interactions (PrePCI) database which comprises over 5 billion predicted interactions between nearly 7 million chemical compounds and 19,797 human proteins. PrePCI relies on a proteome-wide database of structural models based on both traditional modeling techniques and the AlphaFold Protein Structure Database. Sequence and structural similarity-based metrics are established between template proteins in the Protein Data Bank, T, that bind small molecules, C, and proteins in the models database, Q. When these metrics pass a sequence threshold value, it is assumed that C also binds to Q with a probability derived from machine learning. If the relationship is based on structure, this probability is based on a scoring function that measures the extent to which C is compatible with the binding site of Q as described in the LT-scanner algorithm. For every predicted complex derived in this way, chemical similarity based on the Tanimoto Coefficient identifies other small molecules that may bind to Q. A likelihood ratio for the binding of C to Q is obtained from naïve Bayesian statistics. The PrePCI algorithm performs well under different validations. It can be queried by entering a UniProt ID for a protein and obtaining a list of compounds predicted to bind to it along with associated probabilities. Alternatively, entering an identifier for the compound outputs a list of proteins it is predicted to bind. Specific applications of the database are described and a strategy is introduced to use PrePCI as a first step in a docking screen.

## Introduction

Protein-small molecule interactions, termed here protein-compound interactions (PCIs), play essential roles at all biological levels.^1–5^ Delineation of PCIs for the human proteome is essential for developing a systems-level understanding of biological networks and of the molecular basis of therapeutic and off-target effects of drugs. Recent advances in mass spectrometry have enabled high-throughput identification of PCIs for focused sets of metabolites and drugs, uncovering many previously unreported PCIs,^4–7^ suggesting that much of the protein-compound interactome remains to be discovered. We previously reported LT-scanner,^8^ a template-based method which uses protein structural alignment between models of query proteins and experimentally determined protein-compound complexes to predict PCIs involving 26K compounds in the Protein Data Bank (PDB).^9^ Here we describe Predicting Protein Compound Interactions (PrePCI), which extends LT-scanner by dramatically increasing the number of compounds and proteins considered. Calculating chemical similarity with the Tanimoto Coefficient (TC)^10^ between molecular fingerprints of PDB ligands and compounds in the PubChem database^11,12^ expands the chemical space explored to nearly 7 million compounds. In addition, combining protein models from the PDB,^9^ the AlphaFold Protein Structure Database,^13,14^ and our in-house homology database PrePMod^15^ provides almost complete structural coverage of the human proteome. PrePCI provides predictions for ∼5 billion PCIs, and, as reported below, is extensively validated.

Many computational approaches have been developed to predict PCIs. Docking-based methods generate poses of compounds in complex with a protein of interest which are subsequently scored to estimate binding affinities by functions typically based on physical forcefields.^16–20^ Such strategies have enabled structure-based virtual screening of hundreds of millions to billions of small molecules and have discovered novel protein chemotypes.^21,22^ However, the computational costs of pose generation and scoring currently prevent docking methods from being applied at a proteome scale. Ligand-based approaches like Similarity Ensemble Approach^23^ and QSAR based methods,^24^ infer novel PCIs based on the similarity of query compounds to seed compounds that are already known to bind the query protein.^25^ Such methods are able to leverage increasingly large amounts of high-throughput screening data and are generally rapid enough to use with large-scale chemical libraries. However, since most proteins do not have a set of known binders against which to compare, ligand-based methods have difficulty scaling to proteome-wide applications. More recent approaches include 1) machine-learned scoring functions,^26–30^ 2) protein-ligand interaction fingerprints which compare predicted poses to experimentally determined complexes,^31,32^ and 3) convolutional neural networks which learn structural features directly from PCI features and structures.^33–35^

In contrast, proteochemometric methods which infer PCIs using independent protein and compound features are potentially amenable to proteome-scale PCI prediction as they do not require pose generation for each PCI. For example, REMAP, a one-class collaborative filtering approach, formulates PCI prediction as a matrix factorization problem in which low rank matrices representing abstract protein and chemical features are derived from protein sequence similarity and chemical similarity respectively.^36^ 3D-REMAP augments REMAP with ligand binding site similarity and binding affinities for compounds of interest.^37^

LT-scanner uses experimentally resolved protein-compound complexes from the PDB^9^ to scan large databases of protein models to identify residues on their surfaces that are likely to bind similar ligands.^8^ Use of a simple scoring function enables proteome-wide application. The goal of LT-scanner has been to generate novel hypotheses based on protein structural similarity, and this capacity is now enhanced in PrePCI through the addition of protein sequence similarity searches and Tanimoto-based chemical matching. Most closely related to PrePCI is the FINDSITE suite of programs which use protein threading to identify regions within a query protein which can be reasonably modeled using a PDB protein-compound complex as a template.^38,39^ Ligands bound to identified binding sites are then used as seeds for ligand-based virtual screening. Unique to PrePCI is the large proteome-wide database of predicted protein-compound interactions described in this publication.

To illustrate potential applications of the PrePCI database we present two case studies for predictions not detectable by sequence similarity and based on structure similarity and chemical similarity. We anticipate that PrePCI will be a useful resource for generating hypotheses regarding off-target prediction and drug repurposing, lead candidate selection, and identification of metabolites involved in mediating cellular processes. PrePCI predictions can be queried through the web-hosted database application available at https://honiglab.c2b2.columbia.edu/prepci.html.

## Results

### PrePCI overview

The PrePCI algorithm is depicted in Figure 1 and consists of three components. Figure 1A illustrates the sequence similarity component where a query protein sequence is matched to template protein sequences with BLAST. For a given protein-compound pair, the query protein is predicted as a target of a compound which appears in a PDB complex based on the sequence alignment score of the query with the template protein (see Methods). Figure 1B illustrates the LT-scanner component in which a query protein is structurally aligned to a template complex in the PDB^9^ with the Ska program,^40^ which calculates a protein structure distance (PSD) between the aligned structures. Ska emphasizes local versus global alignment with the effect of highlighting structural elements of likely functional relevance, enabling the comparison of the predicted query interface with the template binding site.^40^ The transformation that aligns the two proteins is used to place the PDB compound in the coordinate frame of the query. The LT-scanner scoring function^8^ assesses the compatibility of the compound with the query protein by calculating a score based on the extent to which residues in the query binding site recapitulate the physicochemical interactions between the protein and compound in the template complex. The sequence similarity and LT-scanner calculations are performed for all query protein models and sequences against all PDB protein-compound templates. Figure 1C illustrates the chemical similarity component where PDB compounds are matched to PubChem compounds by TC calculations for their chemical fingerprints. When the TC ≥ 0.5, the compound is predicted to target the protein found by either sequence similarity (Figure 1A) or LT-scanner (Figure 1B). A Bayesian procedure is used to integrate the sequence, structure, and chemical similarity scores for each query protein-compound prediction into a likelihood ratio derived from a true positive set of experimentally characterized PCIs. The scored PCI predictions comprise the PrePCI database (PrePCI/DB).

**Figure 1:**
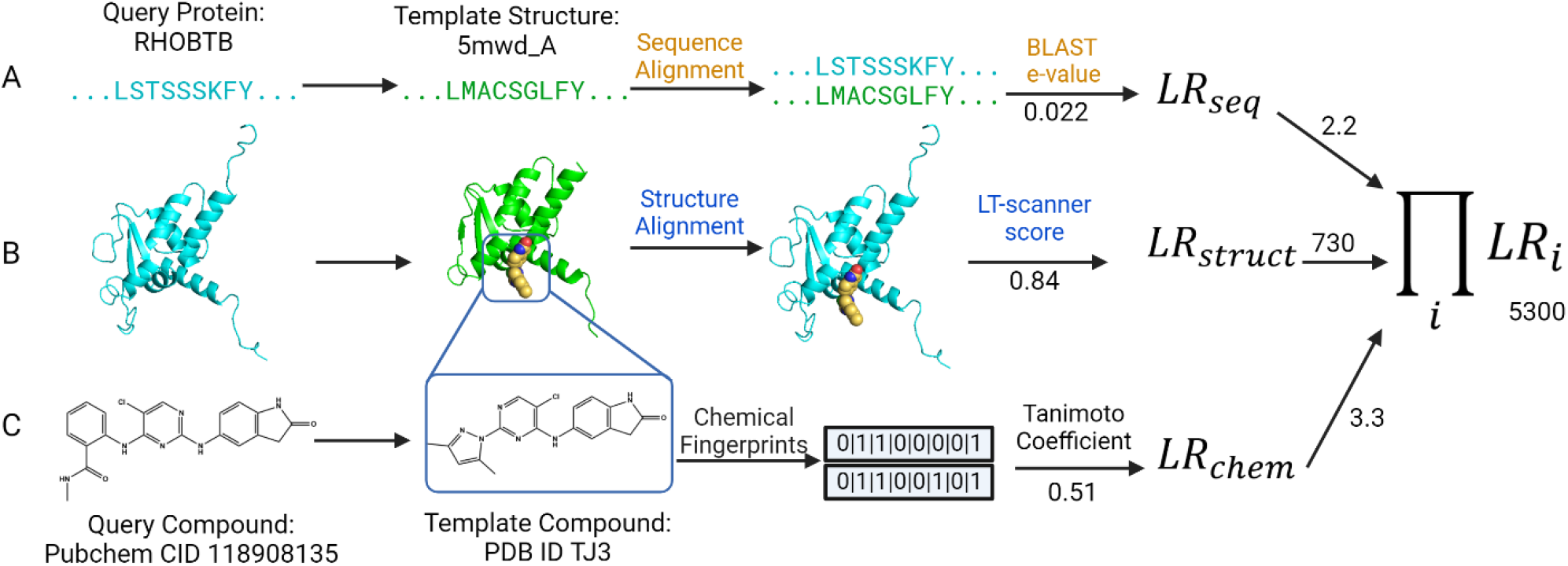
PrePCI algorithm for the prediction of protein-compound interactions. PrePCI uses BLAST and LT-scanner for the query protein (aqua sequence and model, left) to identify proteins within PDB protein-compound complexes that have (A) sequence and/or (B) structure similarity to the query. (C) Compounds predicted to bind the query are identified using a Tanimoto coefficient (TC) chemical similarity search of extended connectivity fingerprints representing the compounds. In this example, the sequence and model for RHOBTB are matched to the PDB complex (PDB ID 5mwd) of the BCL6 BTB-domain bound to PDB compound TJ3 (PubChem CID: 137350052) with BLAST e-value 0.022 and LT-scanner score 0.84, respectively. A query compound from PubChem has TC = 0.51 with the PDB compound. The LR for the interaction between RHOBTB and the query compound (PubChem CID: 137350052) is the product of the LRs from sequence, structure, and chemical similarity scores.

An essential component of LT-scanner is a database of structural models for query proteins for most of the proteins and their constituent domains in the human proteome. To date, we have relied on our PrePMod database of homology models.^15^ Currently, PrePMod contains models for 76,816 protein domains as defined by the Conserved Domain Database (CDD)^41^ where 17,150 human proteins have at least one domain modeled. As described in Methods, a database was constructed of models taken either directly from the AlphaFold Protein Structure Database (AF)^13,14^ or obtained after parsing AF models into CDD domains (AF/CDD).^41^ AF/CDD contains 89,645 domain models covering 20,546 proteins while the union of AF/CDD and PrePMod contains models of 90,308 domains spanning 20,599 proteins. This constitutes a significant (∼15-20%) increase in structural coverage which now includes one representative for essentially every coding gene in the human proteome.

### PrePCI Training and Evaluation

To evaluate PrePCI’s performance, a Naïve Bayes Classifier was trained using 10-fold cross-validation with a true positive set of PCIs that have bioactivity data from PubChem.^11,12^ As described in Methods, the true positive set consists of 285K PCIs for 142K compounds and 2,926 proteins. The negative set consists of 417M hypothetical PCIs between the 142K compounds and 2,926 proteins for which PubChem provides no bioactivity data. For each of the ten folds, PrePCI’s performance was evaluated by ranking predictions by their likelihood ratio (LR) and computing the area under the receiver operator characteristic (ROC) curve (AUROC) and the average precision (or area under the precision-recall curve, AUPRC) (Figure 2). The resulting ROC and Precision-Recall curves are highly concordant, with mean AUROC and average precision of 0.828±0.001 and 0.165±0.002, respectively. It is important to note that, due to the size of the negative set, the training set is heavily imbalanced, and random precision would thus be 7×10^−4^. The average precision of 0.165 therefore constitutes a substantial enrichment of true positive predictions and is likely an underestimate as many PCIs considered false positives likely correspond to as yet undiscovered true interactions.

**Figure 2:**
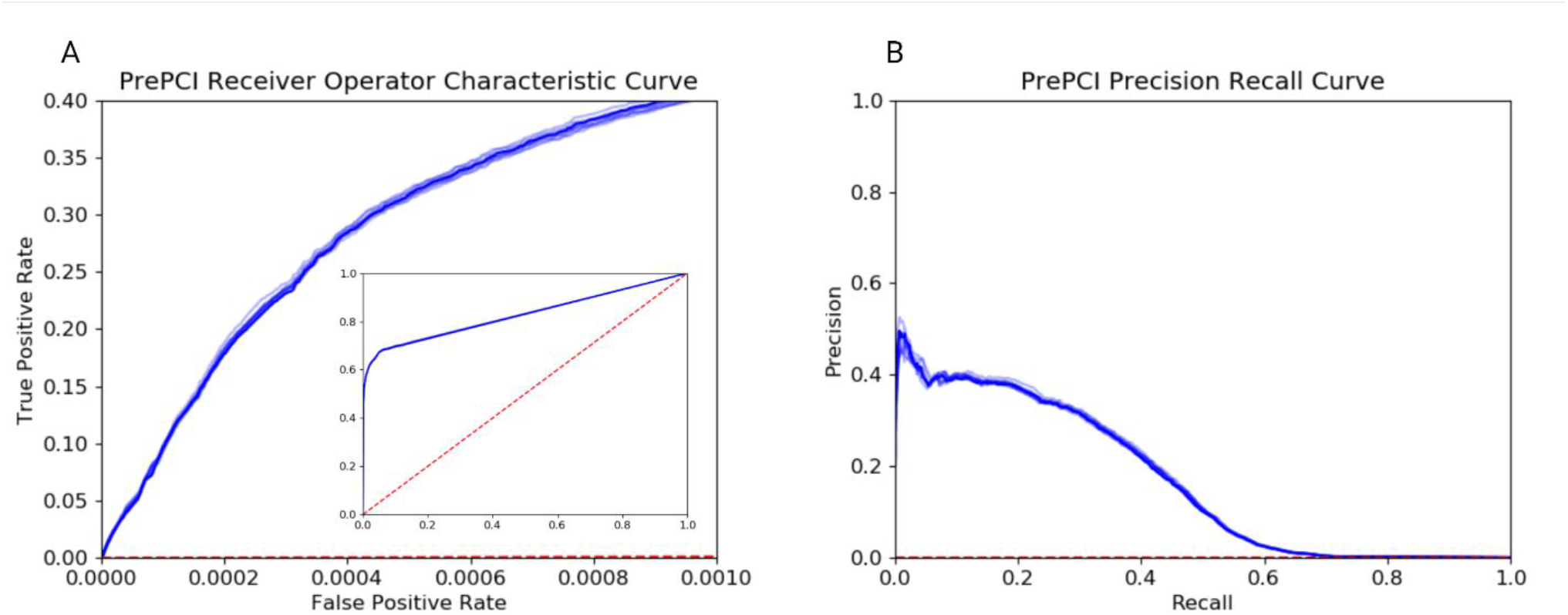
PrePCI performance on unbiased, all-against-all experimental protein-compound data from PubChem. (A) Receiver operating characteristic (ROC) curve and (B) Precision Recall curve for each of the 10 folds of cross-validation for training and testing PrePCI on experimentally observed PCIs from PubChem. Curves for the median area under the ROC curve (AUROC, A) and average precision (B) across all cross-validation folds are darker, while curves for all cross-validation folds (lighter and darker blue) are included to display the range of results obtained from the individual folds. PrePCI’s average AUROC and Average Precision on the PubChem dataset are 0.828±0.001 and 0.165±0.002 respectively.

### LT-Scanner and Sequence Similarity are Synergistic

PrePCI performance was compared to the ability of sequence similarity versus LT-scanner to predict targets by ranking PCIs according to sequence similarity score and LT-scanner score of the template compound, respectively.^8^ Sequence similarity outperforms LT-scanner and performs comparably to PrePCI (Figure S1). Since a sequence identity cutoff between the query and template protein sequences was not implemented, many of the sequence similarity targets are likely to be obvious, but are included in the PrePCI/DB for completeness. In contrast, the unique feature of LT-scanner is its ability to identify non-trivial relationships, for example, in Figure 1 for RHOBTB (see also below for additional examples). Indeed, as can be seen in Table S1, LT-scanner identifies many more relationships than available from sequence similarity alone. The combined use of sequence and structure yields the greatest coverage of true positive PCIs without impairing performance as each method identifies PCIs not detected by the other at comparable LRs (Figure S1).

### The Union of Homology Models and AlphaFold Structures Increases PCI Coverage

PrePCI performance was evaluated with predictions based on PrepMod versus AF/CDD query models. As shown in Figure S2, performance is similar regardless of the query model database used. This reflects the expectation that the LT-scanner score is not strongly dependent on model accuracy. While the number of predictions is greater with AF/CDD versus PrePMod models, the combination of the databases is synergistic and results in the highest number of PCI predictions (Table S2). For example, as depicted in the first row of Table S2, in cases where the query model aligns well with the template complex (LT-scanner score ≥ 0.6), PrePCI-PrePMod predicts 64K PCIs and PrePCI-AF/CDD predicts 77K PCIs. The intersection of the two sets is 39K PCIs whereas the union is 101K PCIs.

### Evaluation with Docking Benchmarks

Two widely used benchmarking datasets, the Directory of Useful Decoys Enhanced (DUD-E)^42^ and the Demanding Evaluation Kits for Objective In Silico Screening 2.0 (DEKOIS 2.0),^43^ were used to further evaluate PrePCI. DUD-E and DEKOIS 2.0 contain PCIs for both active compounds and decoy compounds. DUD-E contains 95 human proteins and over a million PCIs with 19K active compounds and 1.2M decoy compounds. Using leave-one-out cross-validation across the 95 targets (Figure 3A,B, Table S3, Table S5), PrePCI achieves a mean average precision of 0.64±0.20 and an average enrichment factor, EF_0.01_, of 51.0±12.5. The enrichment factor measures how known compounds rank versus a background of decoy compounds, and EF_0.01_ is the enrichment within the top 1% of PCI predictions. DEKOIS 2.0 contains 69 human proteins and 87K PCIs with 2.6K active compounds and 77K decoy compounds. Across the 69 targets (Figure 3A,C Table S4, Table S5), PrePCI achieves an average precision of 0.62±0.19 and an average enrichment factor, EF_0.01_, of 26.6±6.5 on leave-one-out cross-validation.

**Figure 3:**
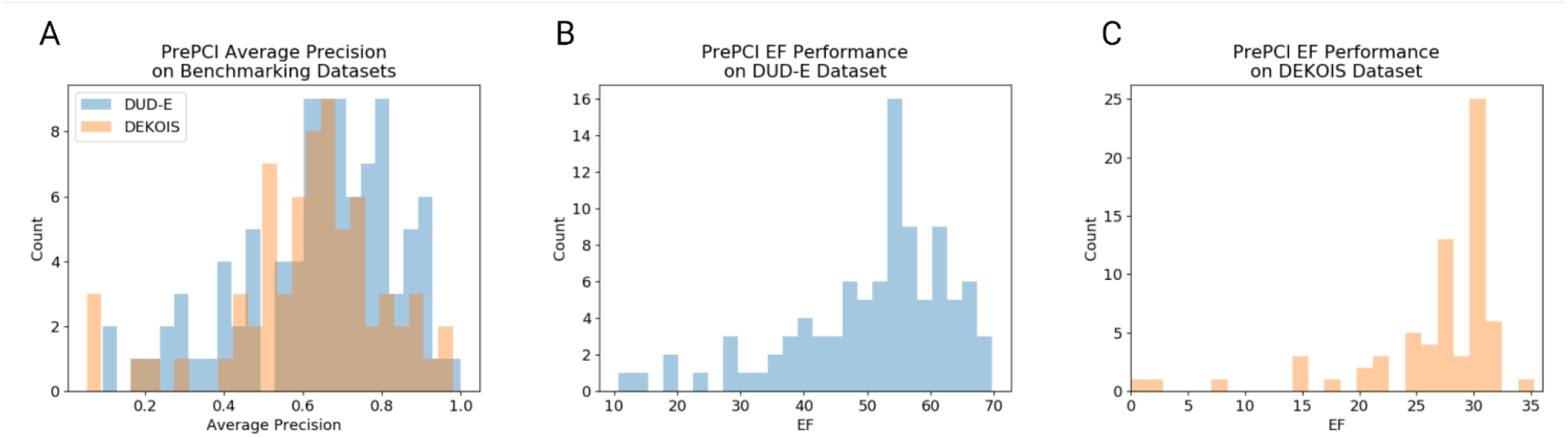
PrePCI performance on DUD-E and DEKOIS 2.0 PCI datasets. Histograms indicate the distribution of (A) per-protein average precision for the DUD-E (blue) and DEKOIS 2.0 (orange) datasets and the distribution of EF_0.01_ for the DUD-E (B) and DEKOIS 2.0 (C) datasets.

The lower average precision value obtained from training with PubChem (0.16) versus DUD-E (0.64) and DEKOIS 2.0 (0.62) (Table S5) is consistent with the expectation that highly imbalanced positive and negative sets lead to an underestimation of true positives. Specifically, the negative set is 1400 times as large as the positive set for PubChem whereas it is typically only 60 and 30 times as large for DUD-E and DEKOIS 2.0, respectively.

### The PrePCI database – PrePCI/DB

PrePCI predictions are available through a web-hosted searchable database (PrePCI/DB) at https://honiglab.c2b2.columbia.edu/prepci.html. PrePCI/DB contains predictions for ∼5 billion PCIs involving 6.8M compounds and 75,643 CDD domains representing 19,797 human proteins. Users can query the database for proteins (with UniProt Accession ID or gene name) or for compounds (PDB compound ID or PubChem ID) to obtain PrePCI predictions for compounds and targets, respectively. Searching by protein will return a list of PDB compounds predicted to bind the protein by either LT-scanner (Figure 1B), sequence similarity (Figure 1A), or both, along with the corresponding LT-scanner scores, BLAST e-values, and LR (Figure 4). The “Click to view PCI” icon will trigger the website to display interactive JSMol windows for exploration of the predicted binding interface as well the structural superposition of the query protein model and the PDB template complex. PDB-formatted files for both the interaction model and the structural superposition can be downloaded for further analysis including more detailed docking studies, as described below. Also on the protein search results page, PubChem compounds similar to the predicted PDB compounds (Figure 1C) can be retrieved by clicking on the “Click to Find Other PCIs” icon which will open a new tab containing all PubChem compounds that are similar to the selected PDB compound (Figure 4). Alternatively, users can query the database for a compound in the form of a PDB ID or PubChem CID. If the compound is present in PrePCI/DB, a list of predicted protein targets is provided, and users can similarly view the interaction models for the protein and template PDB compound and the query-template structure alignments. The following case studies illustrate how both strategies – querying PrePCI/DB for predicted targets and for compounds – can be used to discover novel therapeutically interesting lead compounds.

**Figure 4:**
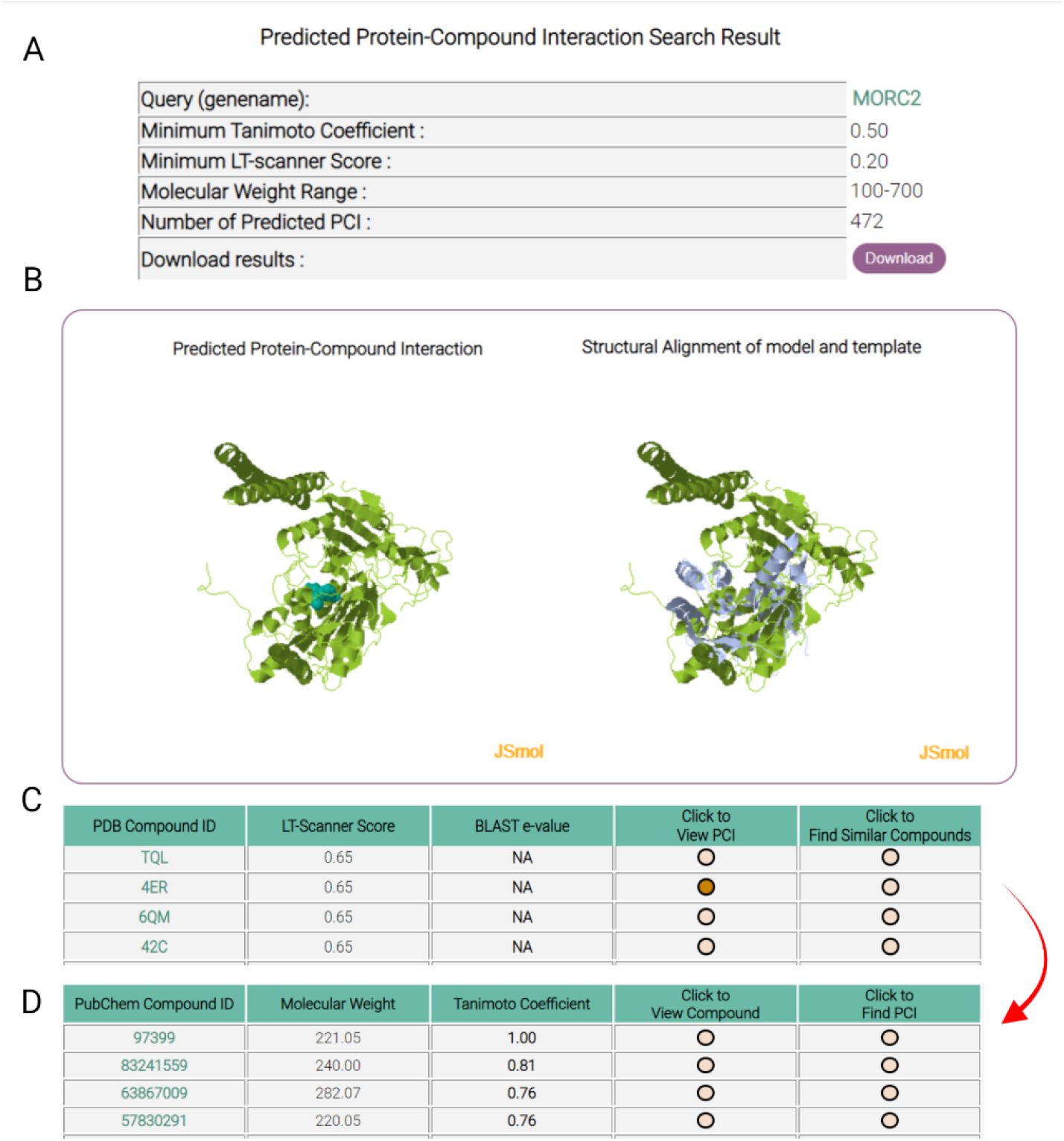
PrePCI webpage output. (A) The top of the webpage displays search criteria and the number of PCIs predicted. (B) Two JSmol windows display the query protein-compound interaction model (left) and query-template superposition (right) which a user may manipulate. (C) A table displays the PDB compounds predicted to interact with the query protein and their corresponding LT-scanner scores and BLAST e-values. The “Click to View PCI” triggers the webpage to display the interaction and superposition models (panel B) while “Click to Find Similar Compounds” opens (D) a new webpage listing PubChem compounds similar to the query PDB compound, which can in turn be used to search for target proteins via the “Click to Find PCI” button.

### Applications of PrePCI/DB

#### Lead fragments for a structure-based docking campaign

One application of PrePCI is to generate a set of lead compounds for focused docking screens. Beginning with a protein of interest, PrePCI/DB provides a set of PDB compounds predicted to bind the query protein (Figures 1 A, B) as well as structurally similar compounds (Figure 1C) that can be screened by docking. Here, we focus specifically on Microrchidia 2 (MORC2), an ATPase involved in epigenetic silencing and transcriptional regulation,^44^ and derive drug-like lead compounds predicted to disrupt its activity. The Hippo signaling pathway has been shown to play critical roles in inhibiting hepatocellular carcinoma (HCC), and the ATPase domain of MORC2 is required for the epigenetic suppression of Hippo signaling.^45^ Therefore, MORC2-ATPase inhibitors, which may serve to reactivate Hippo signaling, have been suggested as a novel approach for HCC therapy.^45^ PrePCI predictions for MORC2 can be leveraged as structural models for more in-depth computational screening and optimization.

PrePCI/DB provides 62 novel compounds with LT-scanner scores ≥ 0.6 for MORC2 (Figure 5A), none of which is predicted by sequence similarity (Figure 1A) and, thus, constitute predictions based solely on structural evidence (Figure 1B). Among the top scoring compounds are the drug-like PDB ligands 4ER (PubChem ID 97399) and 4EU (PubChem ID 73852) (Figure 5A). The LT-scanner models for MORC2/4ER and MORC2/4EU provide a springboard for computational chemical screening of 1,175 PrePCI identified compounds (Figure 1C) with the docking software Glide.^17,18,46^ PubChem compound 139687723 was predicted as the highest affinity ligand with a GlideXP score of −10.0 kcal/mol, which exceeds the predicted affinity with ATP, MORC2’s natural ligand (−9 kcal/mol). A second round of screening (with structurally similar compounds retrieved from PubChem) yielded a compound (PubChem ID 146549823) with a predicted binding affinity of -15 kcal/mol. The docked complex, depicted in Figure 5B, highlights the five hydrogen bonds and the salt bridge between the compound and MORC2, and provides a model for additional pharmacologic optimization.

**Figure 5:**
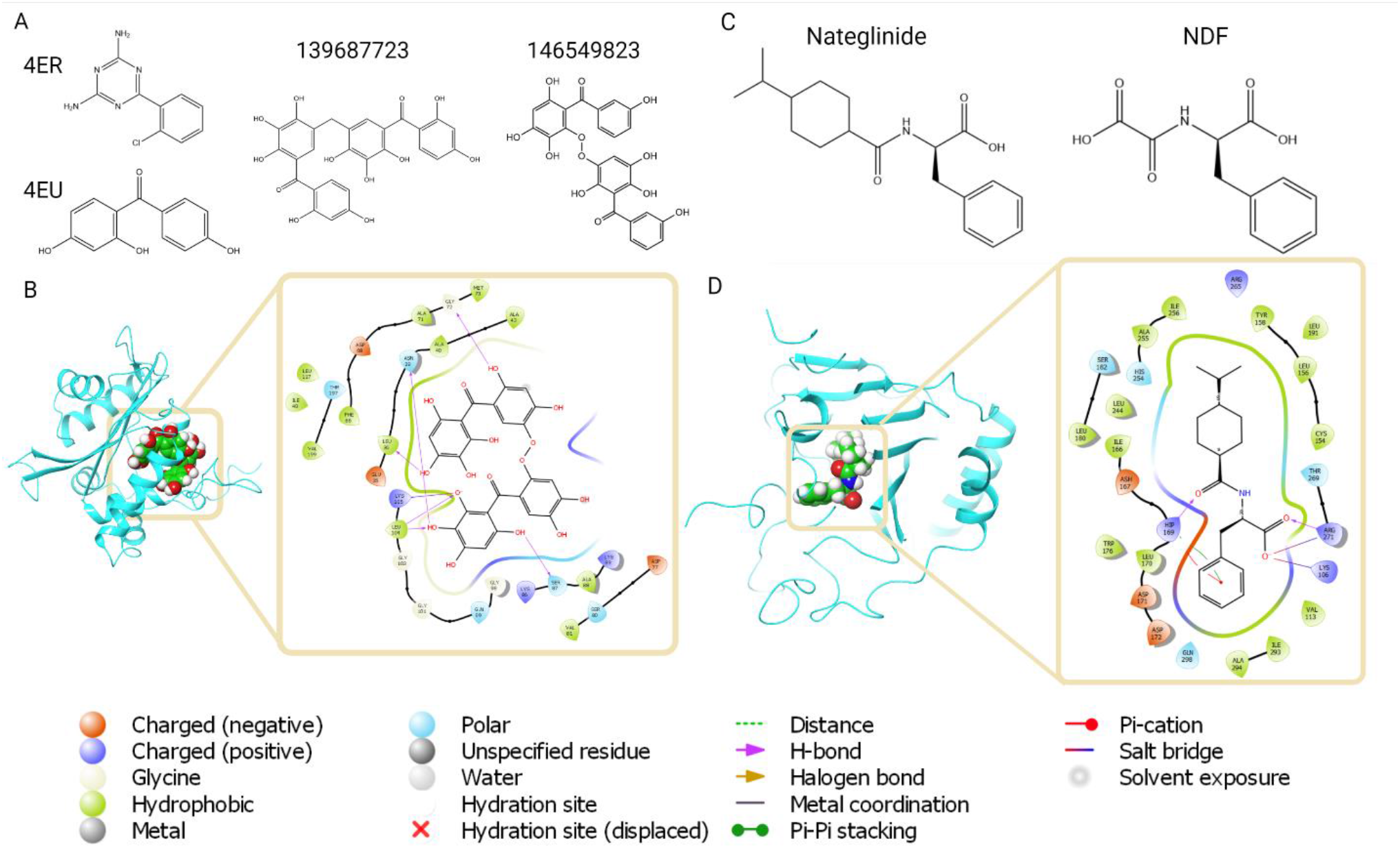
PrePCI guided structure-based virtual screening and drug repurposing. (A) PrePCI predicts 472 PDB compounds bind MORC2, including 4EU and 4ER (left). Chemical similarity and docking predict two PubChem compounds, 139687723 (center) and PubChem compound 146549823 (right), bind MORC2 with high affinity. (B) The docking pose (left) is depicted as a blue backbone ribbon for the target (MORC2) and space-filling representation for the compound (146549823). The diagram (right) highlights atomic interactions predicted by docking. (C) PrePCI predicts 169 proteins bind to nateglinide, including ALKBH4. PrePCI’s prediction that nateglinide (left) binds ALKBH4 is based on its chemical similarity to the PDB compound N-carboxycarbonyl-D-phenylalanine (NDF, right). (D) The docking pose (left) and interaction diagram (right) are shown for the predicted ALKBH4-nateglinide PCI.

#### Novel drug repurposing candidates

Beginning with a compound of interest (Figure 5C), PrePCI can be used to identify potential off-targets of known drugs for repurposing to novel applications. As an example, PrePCI/DB was queried with nateglinide (PubChem CID: 5311309), an anti-diabetic drug, and retrieved ALKBH4, a dioxygenase involved in the demethylation of actin and the promotion of cell migration.^47^ The prediction for ALKBH4/nateglinide arises from the chemical similarity (Figure 1C) between nateglinide and the PDB compound NDF (TC = 0.52; Figure 5C) and the subsequent LT-scanner prediction (score = 0.29, Figure 1B) between ALKBH4 and the template complex HIF1AN/NDF (PDB ID: 1yci_A) – a relationship not detectable with sequence. Moreover, LT-scanner predicts that nateglinide binds directly to key catalytic residues in ALKBH4’s active site^47^ and could therefore act as an inhibitor. Docking simulations with Glide predict a modest affinity of NDF for HIF1AN (score -5.5 kcal/mol) and a stronger affinity of nateglinide for ALKBH4 (score −6.3 kcal/mol) which was increased upon induced-fit docking (score -8.5 kcal/mol; Figure 5D).^48^ ALKBH4 has recently been shown to promote tumorigenesis in non-small cell lung cancer,^49^ and its knockdown induces G1 cell cycle arrest and abrogates proliferation, suggesting repurposing of nateglinide or its optimized derivatives as a treatment for NSCLC.

## Discussion

We have presented the PrePCI algorithm, and a corresponding database PrePCI/DB, which integrates protein structure and chemical and sequence similarity to predict protein compound interactions (Figure 1). PrePCI is an extension of our template-based PCI prediction algorithm, LT-scanner,^8^ which identifies protein targets of small molecules present in the PDB.^9^ The LT-scanner query model database has been updated with structures from the AlphaFold Protein Structure Database providing essentially complete structural coverage for all human protein-coding genes. In addition, PrePCI uses chemical compound similarity based on Tanimoto coefficients among chemical fingerprints to link PDB compounds to compounds in PubChem, which has the effect of increasing the number of compounds that can be explored by more than 200-fold. The increased structural coverage of the human proteome and the expanded chemical space have enabled the prediction of over 5B PCIs, each with component scores and an overall likelihood ratio that allow users to prioritize predictions. Finally, PrePCI achieves high precision for evaluation on curated PCI datasets (PubChem) and docking benchmark datasets (DUD-E and DEKOIS 2.0). More broadly, PrePCI validation on 285K experimentally characterized PCIs involving 142K compounds and 2,926 proteins (Figure 2) strongly indicates that many novel interactions may arise from among the most highly ranked predictions in the negative PCI dataset. We have demonstrated how PrePCI can be used for common medicinal chemistry tasks, such as drug repurposing (Figure 5C,D) and the identification of small chemical fragments as lead compounds for structure-based virtual screening (Figure 5A,B). Importantly, for compounds present in the PDB,^9^ PrePCI generates 3D interaction models and predicts interfacial residues. Given how they are constructed, the models are expected to be crude but can be refined with various docking strategies as illustrated above. While an interaction model is not created for a PCI predicted by chemical similarity (Figure 1C), the predicted compound can be cross-docked to the template PDB compound in the underlying LT-scanner model (Figure 5C,D). In this regard it is important to emphasize that PrePCI is primarily intended for hypothesis generation. PrePCI/DB, which encompasses 5B predicted PCIs involving 6.8M compounds and 19,797 protein targets, is a conveniently accessible structure-informed resource to search for compounds that potentially bind a given protein or, alternatively, proteins that are potential targets of a given compound.

## Materials and Methods

### Template and Model selection

LT-scanner requires a set of query protein structure models and a set of experimentally resolved holo-structures of protein-ligand co-complexes. To select a representative set of template PDB holo-structures, all PDB files in the PDB ligand expo (http://ligand-expo.rcsb.org/) were parsed to identify protein chains bound to ligands and all chains were mapped to their respective UniProt IDs using the SIFTS database.^50,51^ Chains with more than one corresponding UniProt ID, commonly chimeric fusion proteins, as well as chains that did not map to a UniProt ID were excluded. X-ray crystal structures and cryo-EM structures with resolution exceeding 4A and 4.5A, respectively, were removed, and, when a PCI was represented more than once, the highest resolution complex was retained as a template for LT-scanner. This procedure yielded 55,994 unique PCIs between 17,705 proteins and 25,613 compounds after removing compounds with molecular weight < 200Da and fewer than 6 heavy atoms.

### Model Databases

For the human proteome^13,14^ (https://www.uniprot.org/proteomes/UP000005640), structural models for full-length sequences and protein domains as defined by the conserved domain database (CDD)^41^ were constructed as follows.

#### PrePMod

BLAST was used to identify proteins in the PDB with sequences similar to the query sequence. PDB structures identical to the query sequence were retained. For BLAST e-value ≤ 10^−12^, a homology model for the query was created with Nest^52^ with the PDB structure with the lowest e-value used as the template. If no template was identified by BLAST, remote sequence homologs were identified by HHblits^53^ with 5 iterations, and, for e-value ≤ 10^−12^, a homology model for the query was created with Nest^52^ using the PDB structure with the lowest e-value as the template. Otherwise, a homology model for the query was not created. This process yielded PrePMod, a protein model database containing 76,816 domain models spanning 17,150 human proteins.

#### AlphaFold/CDD (AF/CDD)

Models for query proteins were taken from the AlphaFold Protein Structure Database (AF). For proteins with more than 2700 residues, AF provides multiple sequence-redundant models. In these cases, the pLDDT scores (the per-residue confidence metric) were summed across the CDD^41^ domains provided, and the model with the largest total pLDDT was chosen. This procedure yielded 89,645 domain models spanning 20,526 proteins.

Altogether, the combination of PrePMod and AF/CDD provides 166,461 models for 90,308 domains corresponding to 20,599 proteins.

### PrePCI

#### LT-scanner

As described previously,^8^ the structure alignment program ska^40^ identifies templates (T) from PDB protein-compound complexes for a query protein model (Q). T and Q are considered structurally similar when the protein structural distance (PSD) is less than 0.8.^40^ The structural alignment rotates Q into the coordinate frame of T, creating a putative interface between Q and the co-complexed compound (C). Hydrogen atoms are added to T and Q using the Open Babel Package with a pH of 7 and standard pK_a_s.^54^ Four types of interactions between T and C, and Q and C, are identified: 1) hydrogen bonds (distance < 3.5A between heavy atoms and angle > 120), 2) aromatic-aromatic (distance ≤ 5A), 3) ion pairs (distance ≤ 5A), and 4) van der Waals contacts (0.5*r_vdW_ < distance < 1.2*r_vdW_) where r_vdW_ is the sum of the van der Waals radii for the two interacting atoms as defined in the Open Babel parameter set. The extent to which Q is able to recapitulate the intermolecular interactions formed between T and L is calculated by a similarity score, SIM(QL, TL) described previously.^8^ For each potential protein-compound interaction, the LT-scanner score is defined as the maximal observed SIM score between the query protein and any structurally similar template in the holo-structure set. LT-scanner was applied to both the PrePMod and AF/CDD model datasets. In cases where PrePMod and AF/CDD contain models for the same protein/domain, the query model that obtains the higher LT-scanner score is included in the LT-scanner evaluation analyses (Table S2).

#### Sequence similarity

As described previously,^8^ for each of the holo-structure templates used by LT-scanner, BLAST was run using the sequences of each PDB chain as a query against the human reference proteome (UP000005640). For a given PCI, the PDB template containing the compound that yields the lowest e-value with the query protein is identified and assigned an interaction sequence score of –ln(e-value). E-values of 0 were re-assigned to 1e-181, the smallest non-zero e-value obtained from the BLAST results. The sequence similarity component of PrePCI contributes predictions for 17,864 proteins (Table S1).

#### Chemical similarity

Chemical structure data for ∼110 million compounds was obtained from PubChem,^11,12^ and along with the 26K PDB compounds described above, were converted to SMILES format.^55^ Rdkit^56^ was used to compute 1024-bit Morgan2 fingerprints^57^ for each compound, and Tanimoto Coefficients^10^ were computed for each PDB-PubChem compound pair. The reliability of inferring novel compounds from known compounds drops off at Tanimoto Coefficients below 0.5^58^ so only those pairs of compounds with TC ≥ 0.5 were retained, yielding 6,835,528 compounds similar to at least one PDB compound. Overall, PrePCI provides predictions for 6.8M compounds.

#### Naïve Bayes Integration

A Naïve Bayes Classifier was trained to integrate scores into a single likelihood ratio (Figure 1). For each PCI, a reference PDB compound, defined as the PDB compound predicted to bind the query protein by either LT-scanner or sequence that has the highest TC with the query compound, is identified. The chemical, structural, and sequence scores are then defined as 1) the TC between the query compound and the reference compound, 2) the LT-scanner score for the query protein-reference compound pair, and 3) the sequence score between the query protein and the reference compound, respectively. The number of bins was chosen arbitrarily as 10, 10, and 20 for chemical similarity, structural similarity, and sequence similarity, respectively. The scores for each feature were divided into equal intervals and an additional bin was added for NULL scores. Likelihood ratios for each feature and bin were computed as

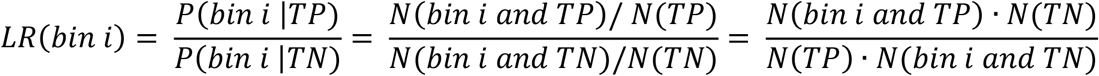

The final likelihood ratio for the PCI is defined as the product of the three component feature likelihood ratios:

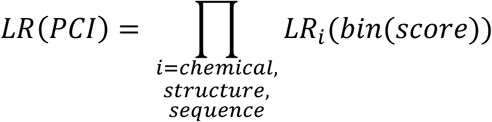

### Experimental Benchmarking

Protein bioactivity data for each protein in the human proteome was downloaded from PubChem’s “Tested Compounds” data from the “Chemicals and Bioactivities Data” section.^11,12^ The data was filtered to retain active, nonredundant experimental PCIs defined as “Active” for the “activity” feature or “<” or “=” for the “acqualifier” feature. This process yielded 1,122,699 PCIs involving 3,559 proteins and 642,498 compounds. Of the 642,498 Pubchem PubChem compounds, 142,490 have Tanimoto Coefficient ≥ 0.5 with at least one PDB compound, and 2,926 of the 3,559 proteins have experimental evidence supporting an interaction with at least one of the 142,490 compounds. After filtering, the true positive set comprised 285,108 PCIs. The true negative set was defined as the 416,640,632 (2926 * 142,490 – 285,108) PCIs that are not present in PubChem. Training was implemented for all pairwise combinations of the 2,926 proteins and 142,490 compounds.

PCI active and decoy compounds for the 102 proteins in the DUD-E database were downloaded from http://dude.docking.org/db/subsets/all/all.tar.gz.^42^ PCI data for the 81 proteins in the DEKOIS 2.0 database was downloaded from http://www.pharmchem.uni-tuebingen.de/dekois/. ^43^ Known binders (actives) and property-matched decoy compounds are provided for each protein in both sets. The true negative sets for the datasets were defined as the provided decoys. Because these datasets contain compounds whose maximal TC with a PDB compound is less than 0.5, 10 additional chemical similarity bins were added to extend the range of TCs down to 0.

The Naïve Bayes classifier was trained using 10-fold cross-validation on the Pubchem Dataset and leave-one-out cross-validation for the proteins in the DUD-E and DEKOIS 2.0 datasets. PCIs with experimentally resolved structures in the PDB were included for training but excluded from test sets to remove recall bias. For each of the cross-validation folds, predictions for PCIs were ranked and ROC and precision-recall curves were created using scikit-learn and matplotlib, and AUROC and average precision were computed using scikit-learn. In addition, the enrichment factor of the top 1% of predictions (EF_0.01_) was calculated to evaluate cutoff-dependent performance:

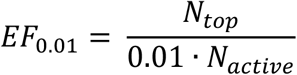

where N_top_ is the number of true positives recovered in the top 1% of predictions and N_active_ is the total number of true positives in the test set.

#### Docking analysis

For all docking studies, a putative interaction model of the PCI was created by aligning the query protein to the template protein using ska.^40^ Proteins structures were then prepared using the Protein Preparation Wizard in Maestro version 13.1 with default settings.^59^ Receptor grids were generated around the query compound setting all neighboring atoms to be rotable and other settings taken as defaults. Ligand structures were prepared using Ligprep; PDB template compounds were prepared from the 3D coordinates within the interaction model predicted by ska and chiralities were inferred from 3D structure whereas 3D coordinates were generated from SMILES strings for non-PDB compounds. Nateglinide was defined using topological SMILES strings to preserve its chirality. For compound screening, multiple conformations were generated for each compound to expand the chemical search space.

#### Docking screen

Structure-based virtual screening was performed to identify novel competitive inhibitors for MORC2. After identifying compounds similar to the PDB compounds 4EU and 4ER, MORC2 and compounds were prepared as described in “Docking analysis.” Similar compounds were screened using rigid-body docking. The standard precision scoring function (SP) was used to rapidly identify promising lead compounds,^60,61^ and the top 25% of predicted compounds were evaluated using the extra precision (XP) scoring function to more rigorously evaluate predicted binding affinity.^17^ This procedure was repeated for compounds similar to the top hit from the first round of screening to identify compounds with higher binding affinity than MORC2’s cognate ligand, ATP.

## Abbreviations and symbols

PCIs: Protein Compound Interactions
PDB: Protein DataBank
TC: Tanimoto Coefficient
AF - AlphaFold, CDD: Conserved Domain Database

## Acknowledgments

This work was supported by the National Institute of Health (grant R35-GM139585, U54-CA209997, BH) (grant T32-GM008224, T32-GM145440, SJT).

## Conflict of Interest

BH is a member of the SAB and consultant for Schrodinger Inc.

## Author contributions

HH, SJT and BH conceived the project, SJT and HH designed the algorithms, SJT carried out the calculations, D Mathur and KB designed the web site, DP provided computational assistance in applications of the models databases, SJT, D Murray and BH analyzed the data and wrote the paper.

## Supplementary Information for

**Figure S1:**
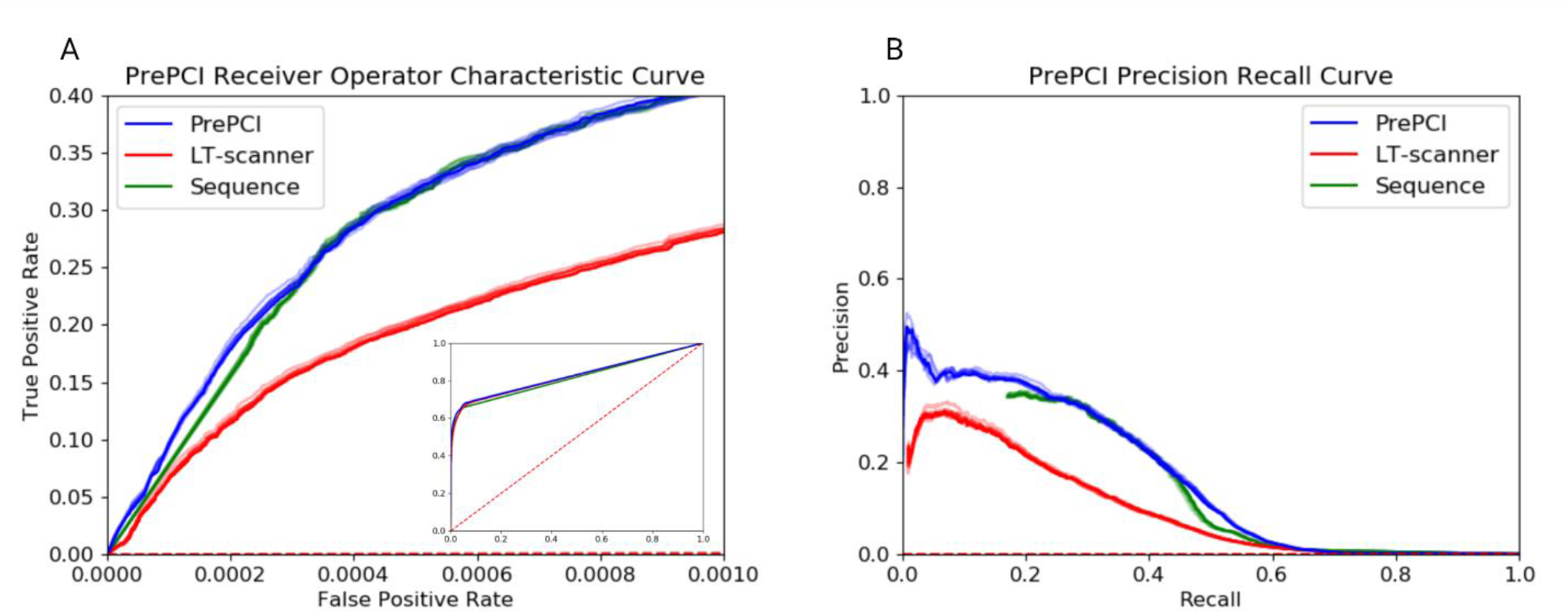
Comparison of the performance of PrePCI, LT-scanner (Structure Similarity), and Sequence Similarity. (A) ROC and (B) Precision-Recall Curves comparing PrePCI (blue) to the performance obtained using LT-scanner and sequence similarity alone, where PCIs are ranked by the LT-scanner score (red line) and sequence similarity score (green line). The average AUROC for PrePCI, LT-scanner and Sequence are 0.828±0.001, 0.825±0.001 and 0.816±0.002 respectively. The mean Average Precision for PrePCI, LT-scanner and Sequence are 0.165±0.002, 0.095±0.001 and 0.150±0.002 respectively.

**Figure S2:**
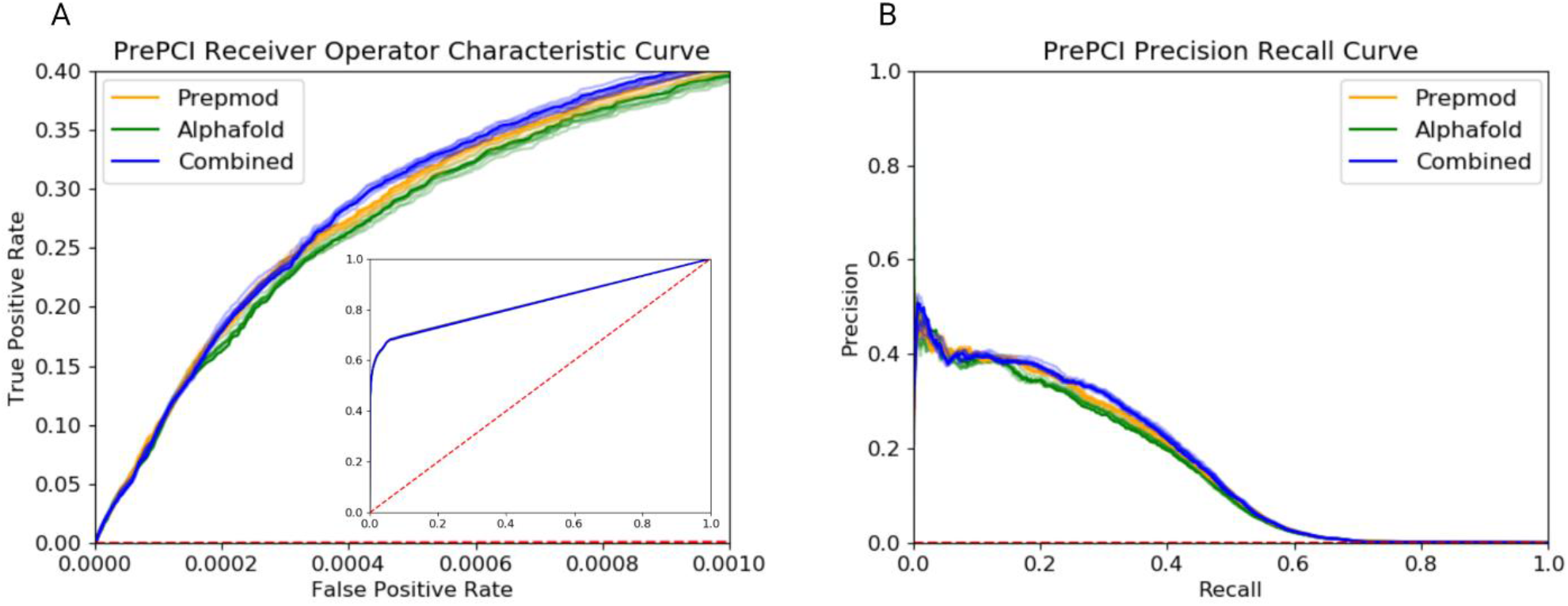
Performance of PrePCI using different model databases. (A) ROC and (B) Precision-Recall curves obtained by 10-fold cross-validation when PrePCI is restricted to structural predictions made using AF/CDD (green) or PrepMod alone (orange) as the model database. For comparison, the results from Figure 2 in which the highest LT-scanner score, irrespective of whether it was based on a PrepMod or AF/CDD model (blue), are included. The average AUROC obtained when using AF/CDD, PrePMod, and the combination are 0.828±0.001, 0.828±0.001 and 0.828±0.001 respectively and the mean Average Precision were 0.163±0.002, 0.156±0.002 and 0.165±0.002 respectively.

**Table S1:** Comparison of the number of PCIs predicted by LT-scanner and sequence-based metrics at comparable LRs. LT-scanner thresholds (Column 1) and Sequence score thresholds (column 2) corresponding to similar LRs were identified. Columns 3 and 4 indicate the number of PCI predictions with LT-scanner scores and sequence scores exceeding their respective threshold. Column 5 indicates the number of PCIs predicted by LT-scanner but not sequence similarity, and conversely, column 7 indicates the number of PCIs predicted by sequence similarity but not LT-scanner. Column 6 indicates the number of PCIs predicted with LT-scanner and sequence scores greater than their respective thresholds (i.e. the intersection of sequence and LT-scanner predictions) while Column 8 indicates the number of PCIs predicted by either LT-scanner or sequence (i.e. the union of the LT-scanner and sequence predictions).

| LT-scanner<br>score<br>threshold<br>(LR) | Sequence<br>Threshold<br>(LR) | LT-scanner<br>PCI<br>Predictions | Sequence<br>PCI<br>Predictions | PCI<br>predictions<br>unique to<br>LT-scanner | PCIs<br>predicted by<br>BOTH LT-scanner and<br>Sequence | PCI<br>predictions<br>unique to<br>Sequence | PCIs<br>predicted by<br>Union of LT-scanner and<br>Sequence |
| --- | --- | --- | --- | --- | --- | --- | --- |
| 0.6 (141) | 186.2791<br>(135) | 101,121 | 62,153 | 72,173 | 28,948 | 33,205 | 134,326 |
| 0.5 (42) | 123.4186<br>(26) | 258,498 | 115,905 | 197,085 | 61,413 | 54,492 | 312,990 |
| 0.4 (10) | 60.55799<br>(10) | 675,105 | 423,786 | 446,875 | 228,230 | 195,556 | 870,661 |
| 0.3 (3.8) | 39.60446<br>(4.0) | 1,566,204 | 749,001 | 1,040,624 | 525,580 | 223,421 | 1,789,625 |
| 0.2 (2.1) | -2.30259<br>(2.2) | 4,449,502 | 2,385,839 | 2,652,348 | 1,797,154 | 588,685 | 5,038,187 |

**Table S2:** Comparison of the number of PCIs predicted by LT-scanner using PrePMod and AF/CDD as model databases. At different LT-Scanner LR thresholds (Column 1), the number of PCI predictions obtained using the model databases indicated are provided. Columns 2 and 3 indicate the number of PCIs predicted by LT-scanner using PrePMod and AF/CDD respectively. Column 4 indicates the number of PCIs predicted when using PrePMod models that are not found with AF/CDD models and conversely, column 6 indicates the number of PCIs detected by AF/CDD and not PrePMod. Column 5 indicates the number of PCIs identified by both PrePMod and AF/CDD models (ie the intersection of PrePMod and AF/CDD predictions) while Column 5 indicates the total number of PCIs predicted by LT-scanner using both model sets (ie the union of PrePMod and AF/CDD predictions).

| LT-scanner threshold | PCIs predicted by PrePMod | PCIs predicted by AF/CDD | PCI Predictions Unique to PrePMod | PCIs predicted by both PrePMod and AF/CDD | PCI Predictions Unique to AF/CDD | Union of LT-Scanner predictions with PrePMod and AF/CDD |
| --- | --- | --- | --- | --- | --- | --- |
| 0.6 | 63,730 | 76,765 | 24,356 | 39,374 | 37,391 | 101,121 |
| 0.5 | 153,660 | 210,225 | 48,273 | 105,387 | 104,838 | 258,498 |
| 0.4 | 401,728 | 561,365 | 113,740 | 287,988 | 273,377 | 675,105 |
| 0.3 | 1,044,492 | 1,313,141 | 253,063 | 791,429 | 521,712 | 1,566,204 |
| 0.2 | 3,043,045 | 3,410,595 | 1,038,907 | 2,004,138 | 1,406,457 | 4,449,502 |

**Table S3:**
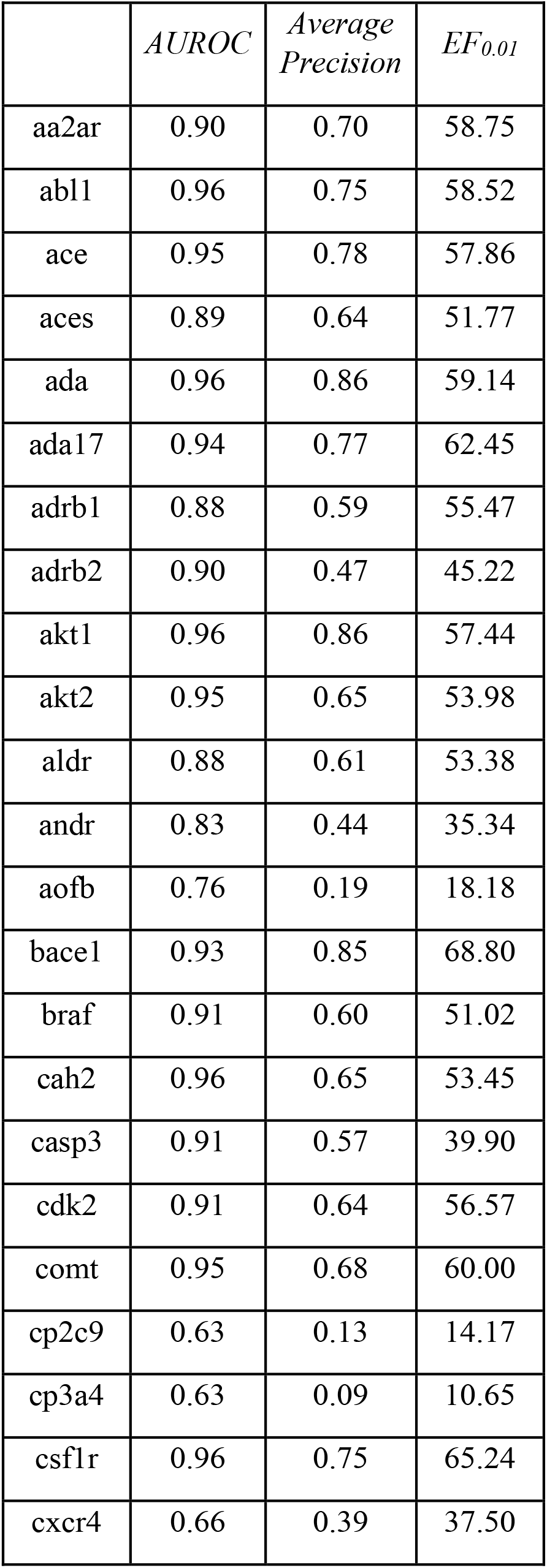
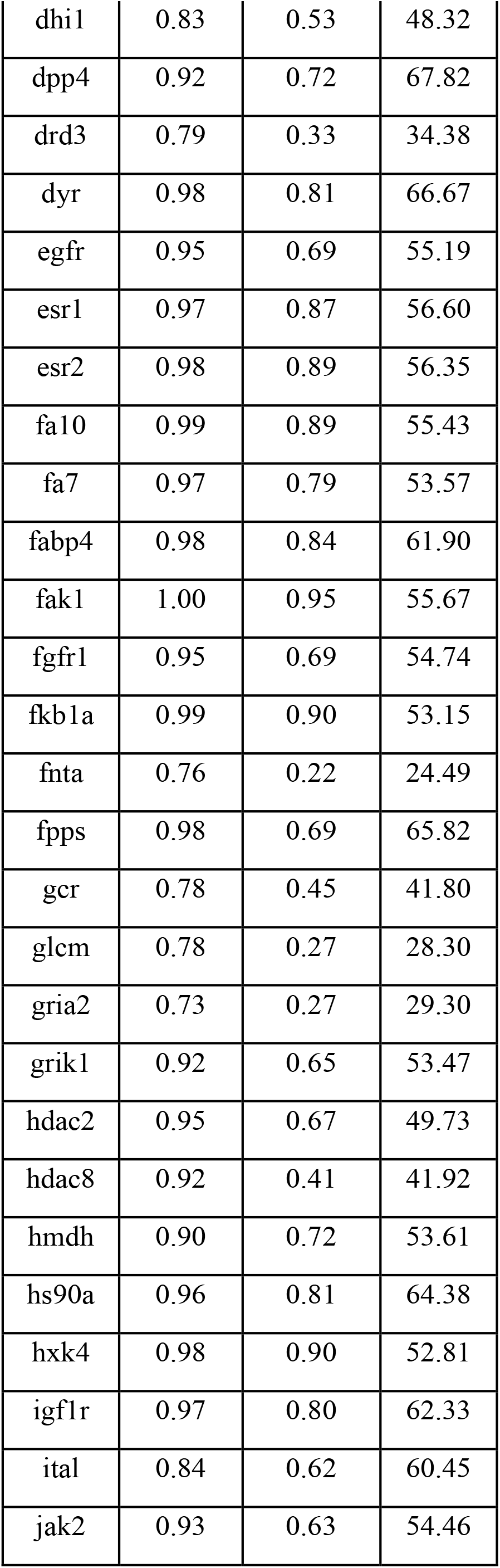
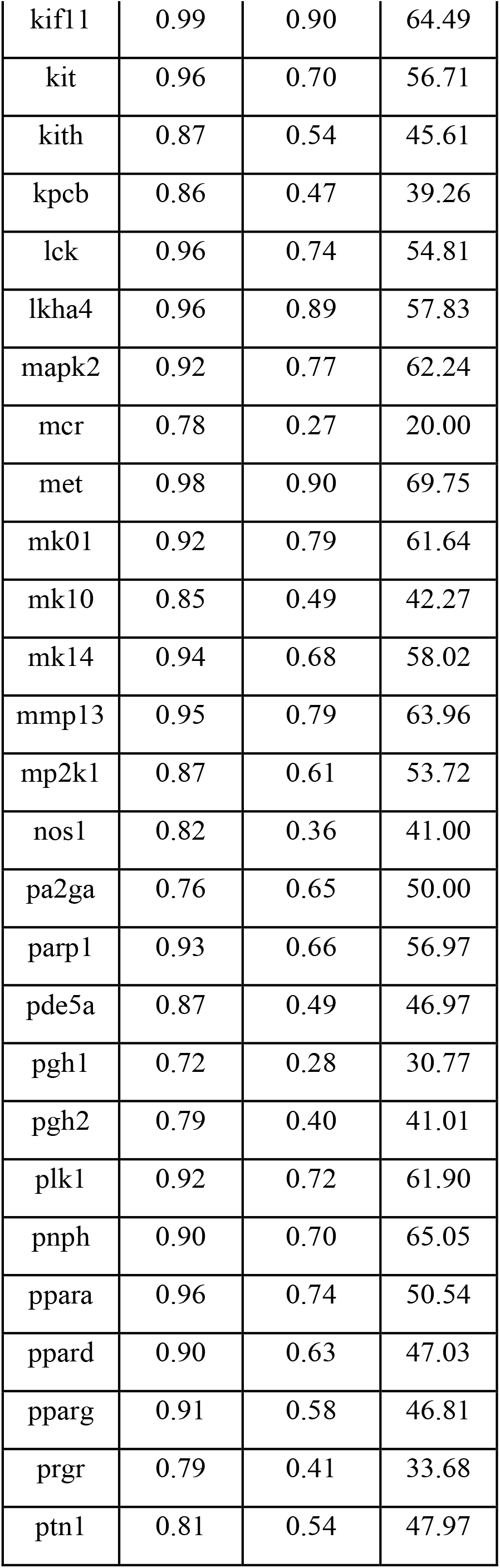
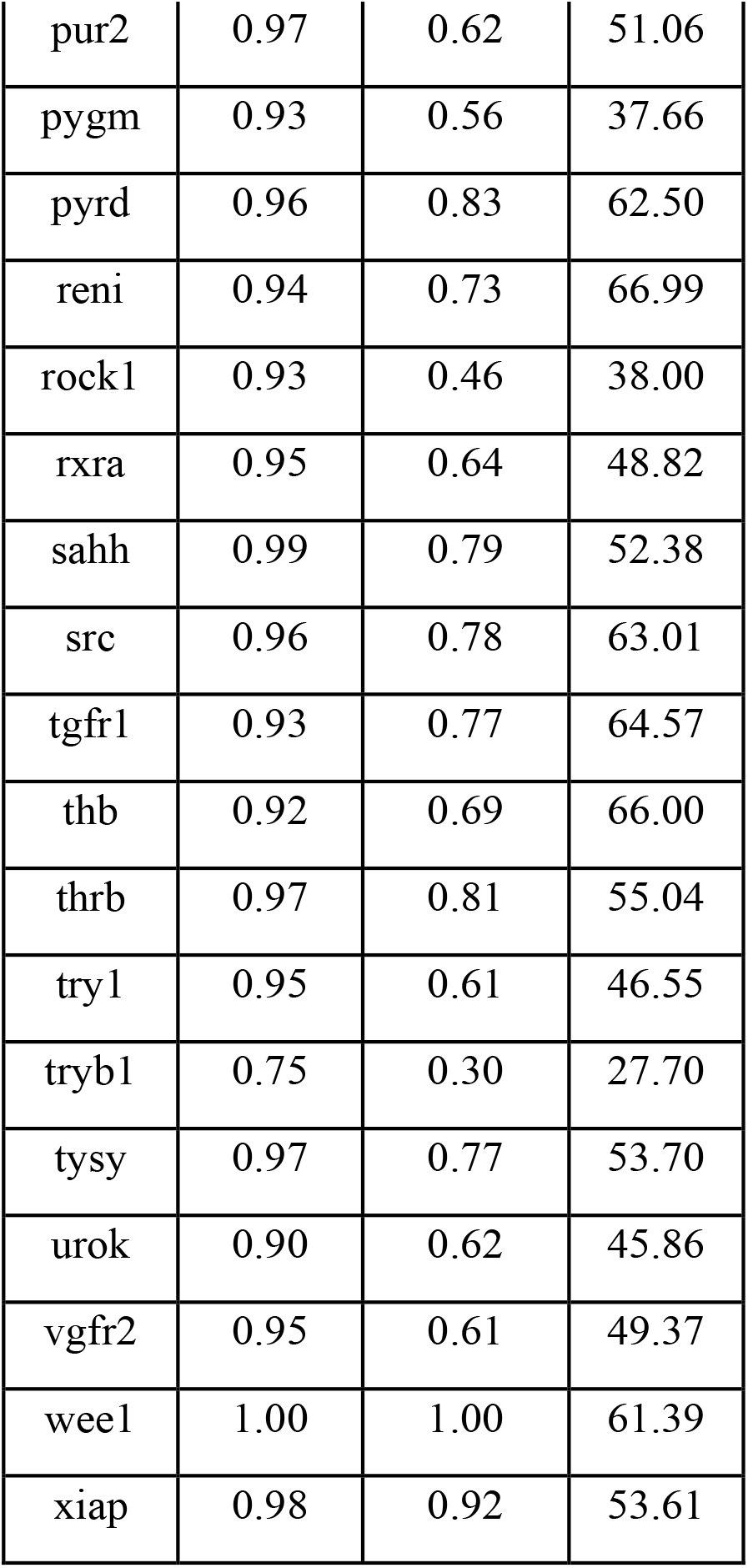
AUROC, Average Precision and EF_0.01_ performance of PrePCI on the DUD-E PCIs.

**Table S4:** EF_0.01_ performance of PrePCI on the DEKOIS 2.0 PCIs.

| *Gene Name* | *AUROC* | *Average Precision* | *EF<sub>0.01</sub>* |
| --- | --- | --- | --- |
| 11betaHSD1 | 0.79 | 0.50 | 30.77 |
| 17betaHSD1 | 0.78 | 0.23 | 15.00 |
| A2A | 0.82 | 0.45 | 25.64 |
| ACE | 0.93 | 0.62 | 25.00 |
| ACE2 | 0.90 | 0.45 | 15.00 |
| ADAM17 | 0.86 | 0.60 | 28.21 |
| ADRB2 | 0.79 | 0.31 | 20.00 |
| AKT1 | 0.89 | 0.68 | 30.00 |
| AR | 0.88 | 0.65 | 30.00 |
| AURKA | 0.93 | 0.70 | 30.56 |
| BCL2 | 0.97 | 0.89 | 30.00 |
| BRAF | 0.93 | 0.66 | 30.00 |
| CATL | 0.93 | 0.74 | 30.00 |
| CDK2 | 0.86 | 0.49 | 28.95 |
| CTSK | 0.99 | 0.74 | 25.00 |
| CYP2A6 | 0.65 | 0.06 | 0.00 |
| DHFR | 0.88 | 0.68 | 30.00 |
| EGFR | 0.85 | 0.50 | 25.00 |
| EPHB4 | 0.96 | 0.65 | 27.03 |
| ERBB2 | 0.99 | 0.82 | 27.50 |
| ER-beta | 0.97 | 0.87 | 30.77 |
| FGFR1 | 0.96 | 0.72 | 27.50 |
| FKBP1A | 1.00 | 0.93 | 30.00 |
| FXA | 0.97 | 0.71 | 27.50 |
| GBA | 0.62 | 0.09 | 7.50 |
| GR | 0.77 | 0.45 | 17.50 |
| GSK3B | 0.90 | 0.58 | 28.95 |
| HDAC2 | 0.97 | 0.81 | 30.00 |
| HDAC8 | 0.96 | 0.78 | 30.77 |
| HMGR | 0.99 | 0.96 | 30.00 |
| HSP90 | 0.97 | 0.90 | 35.29 |
| IGF1R | 0.95 | 0.64 | 20.00 |
| ITK | 0.90 | 0.53 | 27.50 |
| JAK3 | 0.93 | 0.62 | 28.21 |
| JNK1 | 0.90 | 0.64 | 28.95 |
| JNK2 | 0.90 | 0.63 | 30.77 |
| JNK3 | 0.90 | 0.58 | 32.43 |
| KIF11 | 1.00 | 0.98 | 30.77 |
| LCK | 0.93 | 0.63 | 25.00 |
| MDM2 | 0.89 | 0.67 | 30.00 |
| MK2 | 0.92 | 0.75 | 31.58 |
| MMP2 | 0.84 | 0.51 | 25.00 |
| P38-alpha | 0.96 | 0.86 | 30.77 |
| PARP-1 | 0.91 | 0.60 | 31.58 |
| PDE4B | 0.89 | 0.55 | 26.32 |
| PDE5 | 0.87 | 0.54 | 28.21 |
| PDK1 | 0.94 | 0.65 | 31.58 |
| PI3Kg | 0.93 | 0.79 | 32.43 |
| PIM-1 | 0.92 | 0.63 | 29.73 |
| PIM-2 | 0.95 | 0.81 | 30.00 |
| PPARA | 0.92 | 0.60 | 30.00 |
| PPARG | 0.94 | 0.69 | 31.58 |
| PR | 0.79 | 0.39 | 22.50 |
| PRKCQ | 0.74 | 0.20 | 15.00 |
| PYGL-out | 0.99 | 0.86 | 30.00 |
| QPCT | 0.92 | 0.70 | 28.21 |
| ROCK-1 | 0.90 | 0.52 | 22.50 |
| RXR | 0.89 | 0.65 | 30.00 |
| SIRT2 | 0.64 | 0.05 | 2.56 |
| SRC | 0.90 | 0.62 | 27.50 |
| Thrombin | 0.95 | 0.71 | 25.64 |
| TIE2 | 0.97 | 0.68 | 27.50 |
| TK | 0.92 | 0.74 | 30.00 |
| TP | 0.76 | 0.47 | 27.50 |
| TPA | 0.90 | 0.73 | 30.00 |
| TS | 0.89 | 0.56 | 22.50 |
| uPA | 0.84 | 0.58 | 30.77 |
| VEGFR1 | 0.90 | 0.50 | 27.50 |
| VEGFR2 | 0.92 | 0.53 | 26.32 |

**Table S5:** AUROC, Average Precision, and EF_0.01_ of PrePCI performance on the DUD-E and DEKOIS 2.0 datasets.

| DUD-E |  | Mean | Median | Std Dev |
| --- | --- | --- | --- | --- |
|  | AUROC | 0.90 | 0.93 | 0.08 |
|  | Average Precision | 0.64 | 0.67 | 0.20 |
|  | EF | 50.97 | 53.61 | 12.54 |
| DEKOIS 2.0 |  |  |  |  |
|  | AUROC | 0.90 | 0.91 | 0.08 |
|  | Average Precision | 0.62 | 0.64 | 0.19 |
|  | EFs | 26.64 | 28.95 | 6.48 |

